# Clonal propagation history shapes the intra-cultivar genetic diversity in ‘Malbec’ grapevines

**DOI:** 10.1101/2020.10.27.356790

**Authors:** Luciano Calderón, Nuria Mauri, Claudio Muñoz, Pablo Carbonell-Bejerano, Laura Bree, Cristobal Sola, Sebastian Gomez-Talquenca, Carolina Royo, Javier Ibáñez, Jose Miguel Martinez-Zapater, Diego Lijavetzky

## Abstract

Grapevine (*Vitis vinifera* L.) cultivars are clonally propagated to preserve their varietal attributes. However, novel genetic variation still accumulates due to somatic mutations. Aiming to study the potential impact of clonal propagation history on grapevines intra-cultivar genetic diversity, we have focused on ‘Malbec’. This cultivar is appreciated for red wines elaboration, it was originated in Southwestern France and introduced into Argentina during the 1850s. Here, we generated whole-genome resequencing data for four ‘Malbec’ clones with different historical backgrounds. A stringent variant calling procedure was established to identify reliable clonal polymorphisms, additionally corroborated by Sanger sequencing. This analysis retrieved 941 single nucleotide variants (SNVs), occurring among the analyzed clones. Based on a set of validated SNVs, a genotyping experiment was custom-designed to survey ‘Malbec’ genetic diversity. We successfully genotyped 214 samples and identified 14 different clonal genotypes, that clustered into two genetically divergent groups. Group-Ar was driven by clones with a long history of clonal propagation in Argentina, while Group-Fr was driven by clones that have longer remained in Europe. Findings show the ability of such approaches for clonal genotypes identification in grapevines. In particular, we provide evidence on how human actions may have shaped ‘Malbec’ extant genetic diversity pattern.

## Introduction

Clonal propagation is a common practice in perennial crops. In this kind of growing system, a scarce genetic variability could be expected among clones within a given cultivar. However, intrinsic genetic mechanisms such as somatic mutations keep occurring and accumulating along cultivars’ history [^1^]. Grapevine (*Vitis vinifera* L.) cultivars are perennial crops that consist on highly heterozygous genotypes, originated from a sexual cross and clonally propagated to preserve their productive traits [^2^]. Grapevine is among the top five fruit crops in terms of tons produced worldwide [^3^] and it possesses a rather relatively small genome size (~480 Mb) [^4^]. The described features, turn this species into an attractive model for studying the impact of somatic mutations on the genetic diversity of clonal crops [^5–7^]. In this regard, there are many well-documented cases of somatic mutations affecting traits of productive interest in grapevines, mainly involved in berry color determination [^8–10^], berry aroma [^11^], cluster shape [^12,13^] and reproductive development [^14,15^]. However, somatic mutations do not always have qualitative consequences, and quantitative effects have also been reported among clones [^16,17^], even with the responsible mutations identified at the nucleotide resolution level [^18^]. But most of the occurring somatic mutations might not have phenotypic consequences, nonetheless these ‘silent’ variants still constitute a valuable resource of genetic diversity [^5,6^]. For example, they may be used in marker assisted selection programs [^19,20^] or to provide insight on the historical processes shaping current genetic diversity patterns [^1,21,22^].

When analyzing genetic diversity among different grapevine cultivars genetic variation turns very clear [^23,24^]. However, studying the genetic diversity at the intra-cultivar level is more challenging. This limitation is based on the expected low variability and because traditional markers such as SSRs and SNPs selected from inter-cultivar polymorphisms, have shown low efficiency in such approaches [^25–28^]. The increased accessibility to genome-wide scale sequencing has made possible to more accurately address this issue [^5,6,29,30^].

Here we focused on ‘Malbec’ cultivar, which prime name is ‘Cot’ [^31^]. This cultivar has for long been appreciated for the elaboration of high-quality red wines [^32^]. According to the genetic evidence [^33^] and historical records [^32,34^], ‘Malbec’ was originated from the outcrossing of cultivars ‘Prunelard’ and ‘Magdeleine Noir des Charentes’, in Southwestern France (Cahors region). ‘Malbec’ was then introduced into Argentina (Mendoza province) during the 1850s [^32,34^]. In fact, in this South American region is where the largest volumes of ‘Malbec’ wine has been produced for the past two decades [^35^]. ‘Malbec’ shows a notorious clonal phenotypic diversity [^16,36^] and a great adaptation capacity, being successfully introduced into a wide range of agroecological conditions across Argentina [^37^]. However, little is known about ‘Malbec’ inter-clonal genetic diversity. If we track back its clonal propagation history, we can spotlight milestones that could have shaped the current pattern of genetic diversity. Starting from the single seedling that became cultivated after the mentioned outcross, followed by a “bottleneck effect” when it was initially introduced into South America. To the independent accumulation of somatic mutations, as consequence of clonal propagation under different environmental pressures and selection criteria.

In this work, we surveyed ‘Malbec’ intra-cultivar genetic diversity with focus on the impact of its particular clonal propagation history. We implemented a whole genome resequencing (WGR) approach to discover single nucleotide variants (SNVs), occurring among four clones with different historical backgrounds. Then, after a validation process, a reduced set of the identified SNVs was employed to perform a genotyping analysis to survey the genetic diversity across an extensive sampling.

## Results

### a. Genetic diversity among ‘Malbec’ clones is 2000-fold lower than compared to grapevine’s reference genome

We performed WGR of four Malbec clones: MB53, MB59, C225 and C143, that differed in their time span of clonal propagation in Argentina. In total, ~90 million paired-end reads per clone were produced, adding more than 45 Gb of sequence (Supplementary Table S1). Filtered reads were aligned to the *Vitis vinifera* L. reference genome PN40024 [^4^] (hereafter: PN40024), covering ~78% of its length, with a read depth of ~30x (Supplementary Table S1).

After variant calling and filtering processes, we discovered 2,122,796 variants in total (Figure 1). More precisely, we detected 2,121,855 single nucleotide polymorphisms (SNPs), defined here as common variants to the four clones differentiating ‘Malbec’ from PN40024. We also identified 941 single nucleotide variants (SNVs), defined as variants distinguishing ‘Malbec’ clones among each other. From which, 884 were clone-specific (hereafter: CS-SNVs), meaning that one clone had a genotype different from the other three clones. While 57 were shared SNVs (hereafter: Sh-SNVs), meaning that two clones presented the same genotype, different from the other two. Genotypes for CS-SNVs were classified as: Heterozygous (*Het*) = one clone with a heterozygous alternative allele not observed in the other three (253 CS-SNVs); Reference (*Ref*) = one clone showed the reference allele in homozygosis and the other three shared an alternative allele (577 CS-SNVs); and Homozygous (*Hom*): one clone with an homozygous alternative allele and the other three clones were either *Het* or *Ref* (54 CS-SNVs) (Figure 1). Even though CS-SNVs were rather evenly distributed among the four analyzed clones, C143 still showed the highest number (Figure 1). Sh-SNVs genotypes were defined as *Het* and *Hom* when two clones shared the same alternative allele in a heterozygous or homozygous state, respectively. Only a single Sh-SNVs was *Hom*, shared by C143 and C225, that position remained *Het* in MB53 and MB59. The remaining Sh-SNVs were *Het* and distributed as follows: 17 were shared by MB53-MB59 and 17 by C143-C225, while the remaining 22 Sh-SNVs were shared in different combinations: C143-MB53 = 3, C225-MB59 = 4, C225-MB53 = 6 and C143-MB59 = 9.

**Figure 1.**
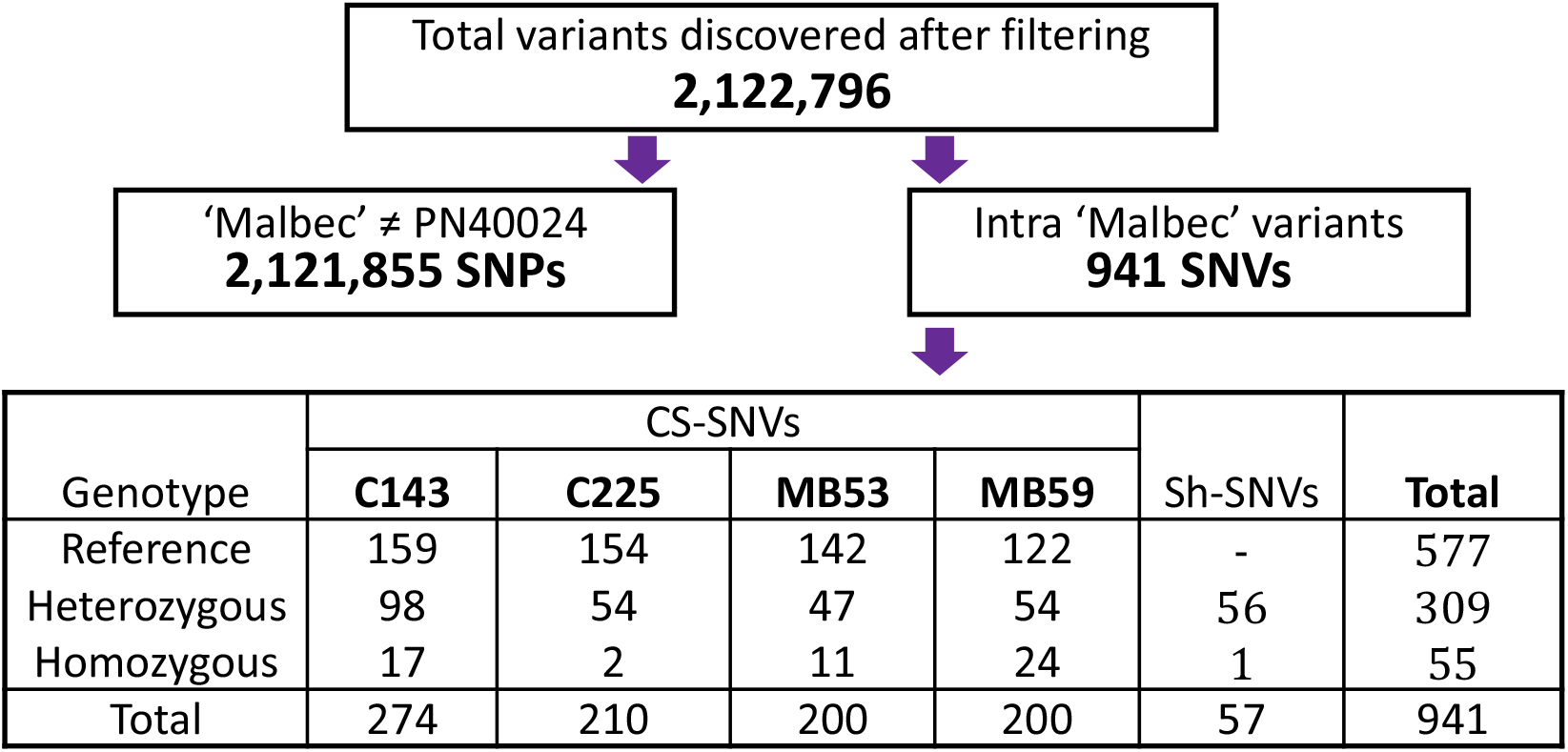
Total single nucleotide polymorphisms (SNPs) and variants (SNVs) identified in ‘Malbec’. SNPs distinguish ‘Malbec’ from PN40024 and SNVs occurred differentially among the four resequenced clones. SNVs are classified based on the clone for which they were identified and also according to their genotype, relative to PN40024. CS-SNVs are diagnostic for variation in a single clone and Sh-SNVs are shared between two clones in different combinations.

We performed a phylogenetic analysis based on the 941 SNVs using PN40024 genotype as an outgroup, and observed that the genetic relations among the four resequenced clones were associated to their clonal propagation history (Figure 2).

**Figure 2.**
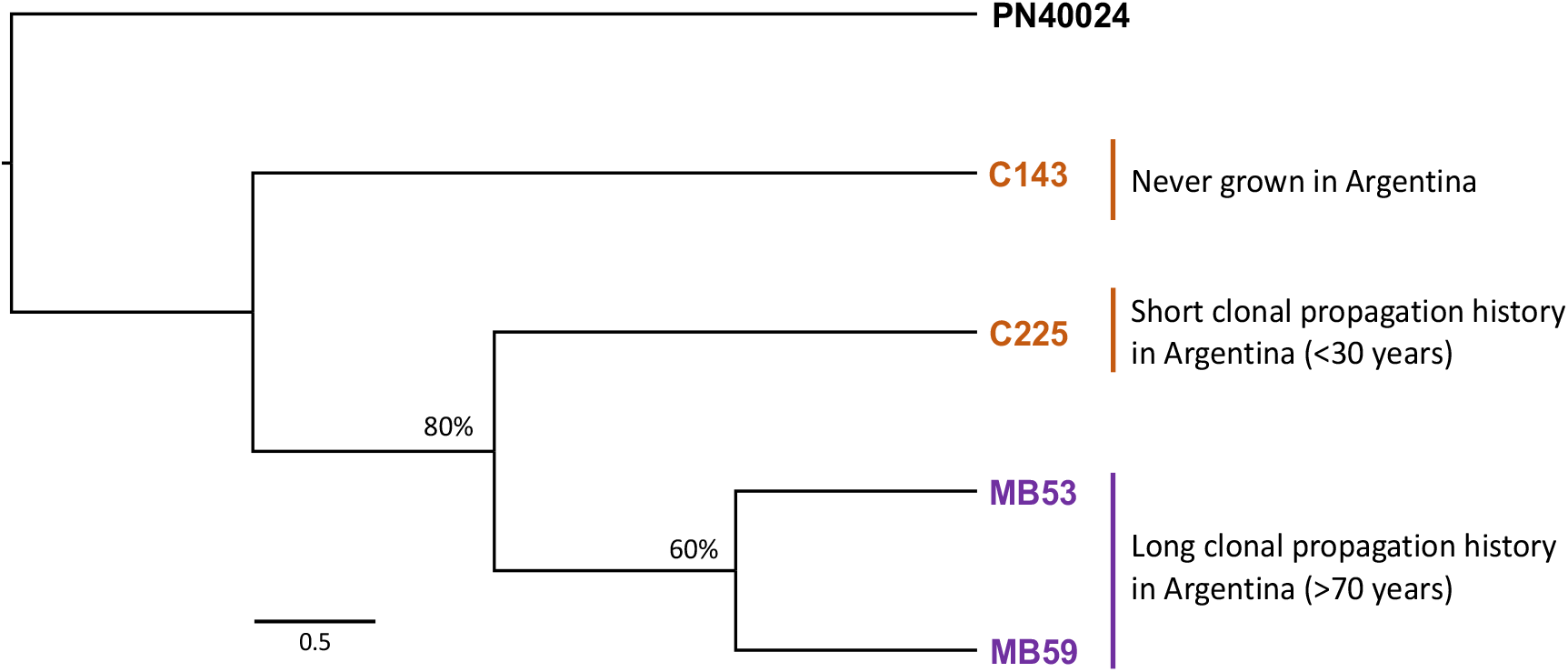
Phylogenetic relations among four resequenced ‘Malbec’ clones. Neighbor-Joining tree based on p-distances and employing 941 SNVs. Percentages on the nodes represent bootstrap supports after 200 iterations, only values >50% are shown. PN40024 genotype was used as outgroup.

More precisely, clone C143 -never grown in Argentina-turned out to be the most genetically divergent from the other three. While C225, with a short history of clonal propagation in Argentina (<30 years), differentiated (80% bootstrap support) from MB53 and MB59. Finally, MB53 and MB59, the two clones that have been longer propagated in Argentina (>70 years) appeared also divergent (60% bootstrap support), but more closely related between each other than to the other two clones.

Out of the 941 described SNVs, 34 were chosen for validation through Sanger sequencing (Supplementary Table S2). All the sequenced SNVs showed the expected allelic states for the corresponding clone, demonstrating the reliability of the employed bioinformatic procedures. As an example, we show the electropherogram alignments of four validated CS-SNVs (one for each of the resequenced clones) (Supplementary Fig. S1).

### b. Genotyping analysis shows that genetic diversity pattern in ‘Malbec’ is related to clones’ propagation history

Genetic diversity was surveyed using a custom-designed genotyping chip. We selected 48 SNVs (including the mentioned 34 validated ones), with 42 CS-SNVs and 6 Sh-SNVs (Supplementary Table S3). Only heterozygous alternative variants were selected, based on their ability to distinguish among the four resequenced clones. Final analyses were performed on 214 successfully genotyped ‘Malbec’ accessions based on 41 properly working SNVs (37 CS-SNVs and 4 Sh-SNVs). We discarded seven out of the 48 starting SNVs and five out of the 219 starting samples, due to technical problems related to missing data. Based on the resequenced clone for which they were originally identified, the 37 properly working CS-SNVs distributed as follows: nine for C143, seven for C225, eleven for MB53 and ten for MB59; while the four Sh-SNVs corresponded to variants shared by MB53 and MB59. Regarding the 41 SNVs variability, as expected for *de novo* mutations, most of them consisted in transitional mutations and only eight were transversion (transitions/transversions=4.1). A total of 22 SNVs markers in the chip were widely informative across the surveyed clonal population, as they ranged from 2 to 164 samples showing the alternative heterozygous allele. Only one of the latter (C225-snv4) showed the three possible genotypes, including the alternative allele in homozygosis. Finally, 19 CS-SNVs showed the alternative heterozygous allele only for one of the four resequenced clones (Supplementary Table S4), which were analyzed with the genotyping chip as a proof of concept of its precision. In fact, the four resequenced clones showed in the chip the expected alternative allele for the respective CS-SNVs, in agreement with the WGR data (Supplementary Table S5).

The genotypes of the 214 samples based on 41 SNVs (Supplementary Table S5), constituted the genotypic dataset used in the subsequent genetic diversity analyses. We built a Median-Joining network, which identified 14 different clonal genotypes: five singletons (i.e. genotypes observed uniquely for one sample) and nine genotypes that were represented by more than one sample (named A to I) (Figure 3). Most genotypes differentiated each other by one, two or three SNVs; except for Genotype-F that accumulated seven, C143 six and MB59 nine SNVs, that differentiated them from their respective closest genotype. The number of samples represented by each genotype ranged from 96 (Genotype-A), comprising 45% of the analyzed accessions, to three (Genotype-I). After inspecting the origin of the samples, no association was observed between the mass selections and the genotypes assignment. Meaning that most genotypes had representatives of samples coming from different mass selections (Supplementary Table S6). The five singleton genotypes corresponded to a sample from *Perdriel* mass selection (Perd_121) and to the four resequenced clones. As expected from the WGR origin of markers in the chip, MB53, MB59, C143 and C225 were the most differentiated samples (Figure 3), due to the effect of CS-SNVs that were variable only for each of them (Supplementary Table S4). Nonetheless, after a more stringent analysis based only on the 22 SNVs showing at least two samples with the alternative allele, the main nine genotypes were recovered (Supplementary Fig. S2). The difference was that C225 and MB53 were the only two samples still differentiating as singletons, while C143, MB59 and Perd_121 were included in Genotypes E, F and A respectively.

**Figure 3.**
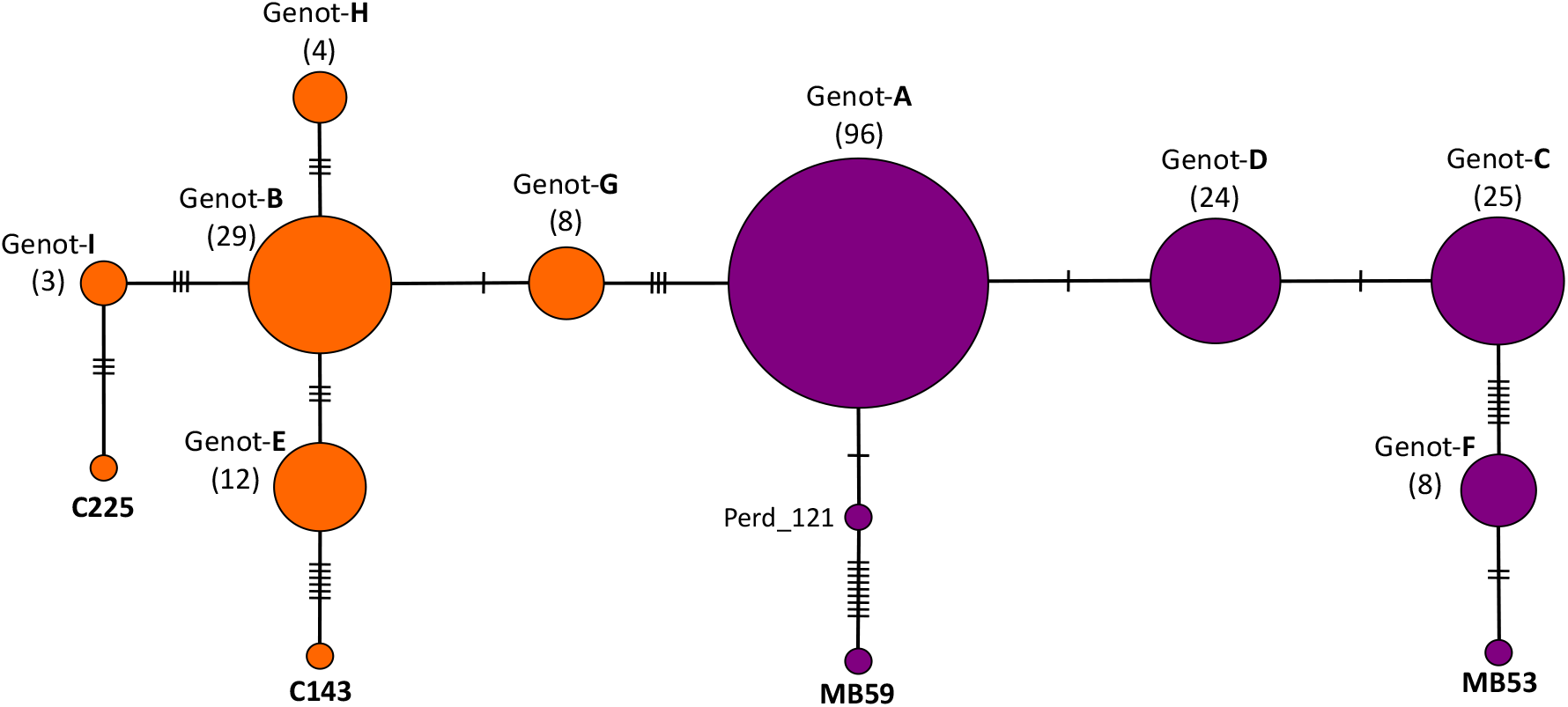
Intra-cultivar clonal genotypic diversity estimated with a custom designed genotyping chip. Median-Joining network was built with the genotypes obtained for 214 clones at 41 SNVs loci. Each circle represents a genotype and its size is proportional to the genotype frequency. In total 14 clonal genotypes were found, nine were represented by multiple samples (named from A to I) and five were singletons (C143, C225, MB53, MB59 and Perd_121). The hashmarks crossing the connecting lines indicate the number of point mutational steps differentiating genotypes. Color code represents Groups Fr (orange) and Ar (purple).

We tested for the phylogenic relations among the 14 identified clonal genotypes based on 41 SNVs. The analysis included a unique sequence representing each of the nine Genotypes from A to I and the five singletons. The resulted tree displayed the existence of two divergent clades, named Group-Ar (Argentina) and Group-Fr (France) (Figure 4a).

**Figure 4.**
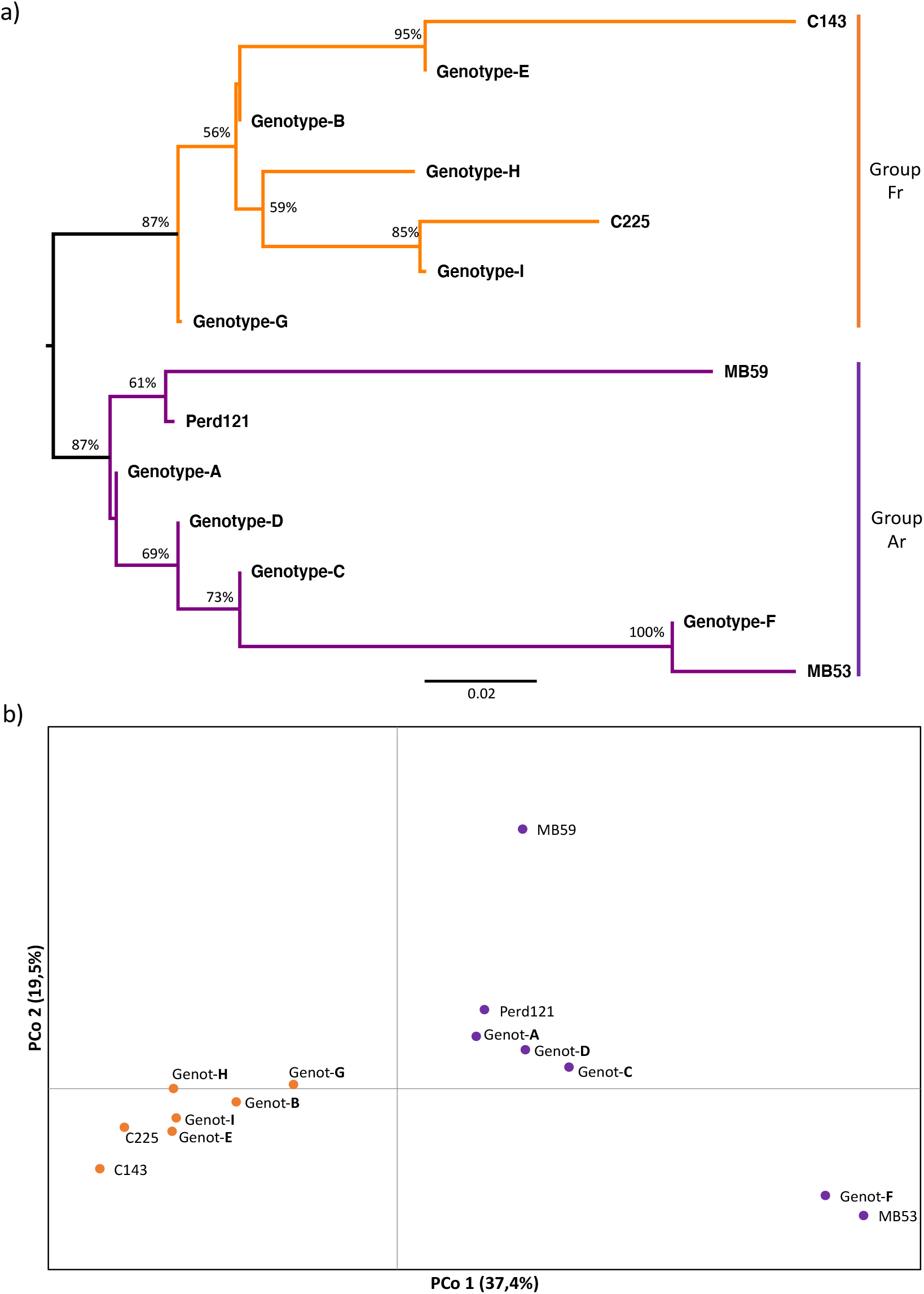
Phylogenetic relationships among the identified clonal genotypes in ‘Malbec’. **(a)** Neighbor-Joining tree based on *p*-distances among the identified genotypes (based on 41 SNVs), nodes bootstrap supports values >50% are shown. The orange clade (Group-Fr) included genotypes closer related to the resequenced clones that have longer remained in Europe (C143 and C225). Purple clade (Group-Ar) included genotypes closer related to the resequenced clones with more than 70 years of clonal propagation in Argentina (MB53 and MB59). **(b)** Principal coordinates analyses (PCoA) based on genetic distances among genotypes. PCoA recovered the same relations than the phylogeny among the identified genotypes, differentiating between Group-Ar (purple dots) and Group-Fr (orange dots).

Group-Ar was driven by the resequenced clones with >70 years of clonal propagation in Argentina (MB53 and MB59) and clustered the closely related genotypes A, C, D and F. Jointly, these genotypes represented the great majority of the analyzed samples (155), including also the singleton genotype Perd_121. While Group-Fr was driven by the resequenced clones that longer remained close to the origin of Malbec in France, never grown or with less than 30 years of clonal propagation in Argentina (C143 and C225), clustering all the genotypes closer related to them: E, B, G, H, I. In total, 64 samples clustered in Group-Fr, including all the other analyzed samples with less than 30 years of clonal propagation in Argentina (Cot42, Cot46, Cot595, Cot596, Cot598, Inta19) or never grown outside Europe (Esp217). Even though Genotype-G was clearly differentiated from other genotypes of Group-Fr (Figure 4a), all performed analyses consistently placed it closer to genotypes from this group (Figure 4b and AMOVA). The distinction between Group-Ar and Group-Fr was also observed in the Principal Coordinate Analysis (PCoA), where the PCo1 and PCo2 explained almost 55% of the genotypic variance (Figure 4b). The separation between the two Groups was mainly depicted by PCo1 (37.4%), all genotypes with a closer genetic distance to C143 and C225 clustered together (including Genotype-G) and the same occurred for genotypes closer related to MB53 and MB59, although with a larger dispersion (Figure 4b). PCoA based only on the four Sh-SNVs, also recovered the distinction between Groups Ar and Fr. Again Genotype-G clearly differentiated from the two Groups, but it was still closer to Group-Fr (Supplementary Fig. S3). Finally, the AMOVA results indicated that a significant proportion of the total molecular variance, was explained after grouping and contrasting the genotypes included in Group-Ar and Group-Fr. The highest AMOVA value was reached when Genotype-G was included in Group-Fr, *Phi*PT = 0,39 (*p* = 0,001).

## Discussion

Extant cultivated grapevines (*V. vinifera* ssp. *sativa*) have retained most of the genetic diversity present in their wild counterpart, ssp. *sylvestris* [^21,38^]. This genetic diversity is evidenced through the great variability observed among cultivars [^7,24^]. However, genetic variation is strongly reduced at the intra-cultivar level. Here, we surveyed the genetic variation in *V. vinifera* L. cv. ‘Malbec’ and found evidence on how clonal propagation history has shaped the diversity pattern of this cultivar.

Somatic mutations mostly accumulate as heterozygous variants, which are more prone to generate false positives in variant calling analyses. Therefore, a major challenge when processing high-throughput genomic data, for clonal genetic diversity studies, consists on avoiding variants overestimation [^39,40^]. Being stringent with the bioinformatic procedures, as well as experimental corroboration of the called variants might provide more certainty on this regard. Here, we worked with a set of variants that were consistently called by three different software: GATK [^41^], BCFTOOLS [^42^] and VARSCAN2 [^43^]. Afterwards, stringent bioinformatic filters, particularly related to the read edit distance and the variant allele frequency (VAF), were applied to discard spurious variants. Moreover, experimental corroboration was successfully performed by means of two alternative technologies, Sanger sequencing (Supplementary Fig. S1) and a Fluidigm genotyping-chip (Supplementary Table S5). All tested SNVs showed the expected alternative allele for the expected sample, so we could assume for a non-significant proportion of false positives among the identified variants. This stringent workflow allowed us to obtain a reliable set of SNVs and set-up a genotyping experiment, to analyze the clonal genetic diversity in ‘Malbec’.

We observed that the number of SNPs distinguishing ‘Malbec’ from the PN40024 grapevine reference genome, exceed the intra-cultivar SNVs by three orders of magnitude (Figure 1). The identified number of SNPs in the present study is within the range of those reported in other works, that have also compared the genetic diversity between grapevine cultivars [^44–46^]. While previous works that studied the intra-cultivar genetic diversity (using WGR data) identified total numbers of SNVs ranging from the few thousand in ‘Chardonnay’[^30^] and ‘Nebbiolo’ [^5^], to the several thousand in ‘Zinfandel’ [^6^]. Here, we present the lowest total number of SNVs reported so far (Figure 1). A reason for this might be that assuming the presence of putative false negatives in the variant calling procedures, we aimed for a stringent filtering to yield reliable markers for clonal lineages identification. However, final results reported in each genetic diversity analysis might be differentially influenced by other technical (e.g. sequencing methods) and biological aspects (e.g. genetic distance among the analyzed clones), as well as by the aim of the analysis [^47,48^]. Regardless of the differences in the absolute numbers of SNVs identified, it is clear that in grapevines the intra-cultivar genetic diversity drops drastically when compared to the inter-cultivar. This observation corroborates the role of vegetative propagation in preserving the desired phenotypes of cultivars, by stabilizing the accumulation rate of novel genetic variation [^2^].

Despite the scarce intra-cultivar genetic diversity in grapevines, the identified variants successfully distinguished 14 different clonal genotypes of ‘Malbec’ (Figure 3). We found no association between the genotype assignment and the mass selection origin of our samples. Plants from the same mass selection share particular phenotypic traits of productive interest (Supplementary Table S7). However, the sought phenotypic homogeneity contrasts with the observed genetic diversity, suggesting that SNVs analyzed here are not associated to genes responsible for the selected traits. On the other hand, the number of samples represented by each genotype was highly variable. Genotype-A was the most abundant, including almost half of the studied accessions (Figure 3). Genotype-A abundancy could indicate that this has been the most propagated lineage in Argentina. Either as consequence of a “bottleneck effect” caused by ancestral introductions of ‘Malbec’ in South America, and/or posterior selections favored by its productive performance. However, we cannot rule out that Genotype-A abundancy could be consequence of a sampling bias. It is expectable that including more samples with diverse origins, as well as employing additional SNVs markers, would turn into a greater number of genotypes represented by fewer samples. Despite of these caveats, even with a reduced set of the identified genetic markers here, it was possible to recover the main identified clonal genotypes (Supplementary Fig. S2).

The identified clonal genotypes clustered in two genetically divergent groups, Groups Ar and Fr (Figures 4a and 4b). The observed pattern of genetic diversity in ‘Malbec’ is likely resembling the combination of natural and human directed processes. The only natural source of genetic variation in grapevine cultivars are somatic mutations and epimutations, which arise during the vine growth and might be pass to daughter vines through vegetative propagation [^49,50^]. Therefore, shared mutated positions turn into fingerprints that provide information on the history of a given clonal lineage [^51^]. Here, by using only four Sh-SNVs was enough to recover the distinction between the two main identified groups (Supplementary Fig. S3). On the other hand, as a species of commercial interest, human actions such as plants transportation, as well as clonal and mass selection are over imposed to the observed patterns of genetic diversity [^2,22,49^].

Historical records report that the first ‘Malbec’ plants were introduced from France to Argentina (Mendoza province) in the 1850s [^32,34^]. After that, wine-producers kept introducing plants into Argentina at a continuous rate, that was slightly increased during the 1990s [^52^]. The found genetic diversity pattern could be reflecting this history. Distinguishing among genotypes that have gone through alternative pathways and accumulated different somatic mutations, and bringing together those with a more recent shared history. Genotypes included in Group-Fr are closely related to the resequenced clones that have longer remained in Europe (C143 and C225), including also all the other analyzed samples that were never grown or were recently introduced into Argentina. On the other hand, genotypes from Group-Ar are closely related to the resequenced clones with a longer time span of clonal propagation in Argentina (MB53 and MB59), suggesting a closer link to those first plants introduced from France. Among the analyzed samples, we can pinpoint those accessions never grown or more recently introduced into Argentina (<30 years). However, we cannot tell with accuracy the exact time span of clonal propagation for those accessions that have remained for more than 70 years in Argentina. In particular, some of the latter accessions appeared included in Group-Fr (Supplementary Table S6). This could be resembling intermediate times of introduction for certain accessions (as those from Genotype-G) or possible traceability inconsistencies. In the same direction, it is not possible to track back the precise history of individual plants from the sampled mass selections; the only available information relates to their vineyard of origin and productive criteria of selection. In this context, it is important to highlight that we were able to retrieve the phylogenetic relations among the four resequenced clones with a known history, by means of the custom-designed genotyping chip.

The set of markers included in the chip proved useful for clonal genotypes distinction. In fact, by genotyping as few as four Sh-SNVs would be enough to tell if a ‘Malbec’ plant is closely related, either to ancestors that were early introduced in Argentina or that longer remained in Europe. At the same time, CS-SNVs were essential at discovering the main clonal genotypes identified here. This observation further supports the importance of combining clone specific and shared variants, to enhance genotyping experiments sensitivity for clonal diversity studies [^30,51^]. Moreover, custom-designed genotyping for grapevine cultivars has already been proven as a valuable tool with different applications. For example, for nurseries to fill with genetic evidence the historical gaps of clonal accessions [^51^], and for the wine industry for traceability and authentication purposes [^53^].

In conclusion, we could setup an efficient workflow to identify a reliable set of clonal genetic variants, that were employed to design an informative genotyping experiment. We were able to distinguish several clonal genotypes within ‘Malbec’ and observed that clonal propagation history has shaped its genetic diversity pattern. Findings add further evidence on the importance of high-throughput genotyping in grapevines as baseline information, to better understand cultivars’ history and as a tool with industrial application.

## Materials & Methods

### a. Biological material

To perform WGR, we obtained young leaves and shoot tips from four ‘Malbec’ clones. Two clones were sampled at the Mercier Argentina nursery collection (Perdriel, Lujan de Cuyo, Mendoza): Malbec-501 (MB53) and Cot-ENTAV-598 (C225). One clone was sampled at Mercier Argentina nursery Granata vineyards (Perdriel, Lujan de Cuyo, Mendoza), Malbec-059 (MB59). The fourth clone: Cot-143 (C143) was sampled at the “*Finca El Encín*” ampelographic collection (ESP-080, Alcala de Henares, Spain). The two accessions labeled as ‘Malbec’ (MB53 and MB59) represent plants with long history of clonal propagation in Argentina, meaning that they have been propagated in this country for more than 70 years (Mercier nursery records). We also included two accessions labeled as ‘Cot’ (C225 and C143), with short and null histories of clonal propagation in Argentina. More precisely, C225 was introduced into Argentina from France (ENTAV-INRA) during the 1990s (Mercier nursery records) and C143 was sampled in a Spanish germplasm collection, therefore it was never grown in Argentina.

For the genotyping analysis, shoot tips and young leaves were obtained from 219 plants. We sampled 70 ‘Malbec’ clonal accessions (including the four resequenced ones) belonging to: (a) the National Institute of Agricultural Technology (INTA-Mendoza) collection (28 clones), (b) Mercier Argentina Nursery collection (37 clones) (c) Mercier Granata vineyard (three clones) and (d) Finca El Encin (two clones). Time span of clonal propagation in Argentina were obtained from [^52^] and from Mercier nursery records. We also obtained 30 samples from each of five different Mercier’s mass selections (150 samples in total), located at Granata vineyards (Perdriel, Lujan de Cuyo, Mendoza). Further details about mass selections and samples are in Supplementary Tables S7 and S8 respectively.

### b. Whole genome resequencing, variant calling and validation

#### DNA extractions and resequencing

Whole genomic DNA from the four ‘Malbec’ clones: MB53, MB59, C225 and C143 was isolated using the DNeasy Plant Mini Kit (Qiagen), including a RNase treatment, according to manufacturer recommendations. DNA quantification and quality checks were performed with NanoDrop 2000 spectrophotometer and agarose gel (5%) electrophoresis. Library preparation and sequencing was performed at the Center for Genomic Regulation (Barcelona, Spain) 125 bp length paired-end reads were produced using the HiSeq 2000 Illumina technology with the Sequencing v4 chemistry.

#### Reads alignment, variant calling and filtering

Standard quality checks of the FASTQ files were performed with FastQC [^54^]. Raw reads were pre-processed following the GATK Best Practices workflow with the toolkit GenomeAnalysisTK-3.3-0 [^41^]. After marking Illumina adapters with Picard toolkit v2.9.4 [^55^], sequences were aligned to *Vitis vinifera* L. reference genome PN40024 [^4^]. We employed the Burrows-Wheeler algorithm as implemented in BWA-MEM v0.7.12-r1039 [^56^], to align our reads to the reference genome. Mapped reads were thoroughly filtered also with Picard toolkit [^55^] allowing only non-duplicates, unique and concordant alignments with a maximum read edit distance of 1 per 25 nucleotides of query sequence [^57^]. Filtered alignments were used as input for variant calling, comparing to PN40024, using three different tools with default parameters and in the multi-allelic mode: GATK UnifiedGenotyper [^41^], BCFTOOLS call v1.9 [^42^] and VARSCAN2 mpileup2cns v2.3.9 [^43^]. Produced gVCF files for each accession were intersected and only those single nucleotide variants identified by all three callers were retained, while INDELs and structural variations were not considered in this study. Bioinformatic procedures were adjusted using a set of SNPs between ‘Malbec’ and PN40024 retrieved from Vitis18kSNP array results [^58^]. Only confident identified raw variants were retained, based on WGR recommendations of total depth (DP), variant allele frequency (VAF), strand bias and distance bias (Bentley et al. 2008). Cut-off values for these parameters were: DP = [15-150]; VAF(Ref) ≤ 0.025; VAF(Het) = [0.25-0.75]; VAF(Hom) ≥ [0.95]; P-value (strand bias) ≤ 0.0001 and P-value (distance bias) ≤ 0.0001. Variant allele frequency ranges were particularly adjusted to reduce -at the minimum possible-the presence of spurious variants. Chimeric mutations are frequent in grapevines, occurring differentially between the L1 and L2 cell layers of the developmental tissue from the apical meristem [^49^]. L1 layer gives rise to the epidermis and represent a smaller proportion of the total tissues conforming a plant (nearly 30%) [^50^]. With the employed VAF filters we expected to detect most of the chimeric mutations occurring in the L2 cell layer, VAF around 0.3 (half of the total frequency in 60% of somatic tissues). While chimeric heterozygous mutations occurring only in the L1 would be mostly excluded. We assumed that variants loss as a trade-off, in the aim of reducing the false positives.

#### Corroboration of the bioinformatic pipeline

We employed IGV v2.3.97 [^59^] to manually corroborate a sub-set of the identified SNVs and to isolate a ~600 bp length sequence containing the target SNVs in the mid-region. These sequences were used as templates for primer design to perform PCRs and Sanger sequencing of the amplicons. In order to avoid both, primer annealing and later genotyping issues, we checked for the absence of variable sites on the 5’- and 3’-regions of the sequence and in the proximities of the SNVs target position. Primers were designed using the Primer BLAST tool [^60^], with an average annealing temperature (Tm) of 60.3°C (range: 58.8-62.5°C) and an average amplicon length of 447 bp (range: 300-582 bp), more details in Supplementary Table S2. PCRs were conducted in a 25 μl final reaction volume containing: 0,3 ul (5 U/μl) Taq Polymerase High fidelity (TransTaq); 1,25 ul (10x) Buffer GC-enhancer (TransTaq); (2,5 ul) 10x PCR Buffer I (TransTaq); 1 ul (2.5 mM) dNTPs; 1 ul of each (10 μM) Primer forward and reverse and 3 ul (40 ng/μl) DNA template. Cycles consisted in a denaturation step of 5’ at 98°C; 35 cycles of 30” 94°C, 30” at 60°C and 30” at 72°C, and final extension of 7’ at 72°C. PCR products were purified using ExoSAP-IT PCR Product Cleanup (Thermofisher), following the manufacturer recommendations. To validate the target SNVs, electropherograms of the four resequenced clones were aligned and inspected with CODON CODE ALIGNER v4.0.4 (CodonCode Corp. USA). SNVs were considered as validated if the allelic state at the position of interest in the sequence, coincided with that observed in the *vcf* file and in the IGV genome browser. For example, for a heterozygous CS-SNVs the clone for which the variant was identified must be heterozygous and for the other three clones must be homozygous as the reference genotype (e.g. Supplementary Fig. S1).

### c. Genotyping

DNA extractions were performed employing the NucleoSpin® Plant II Plant Mini kit (Macherey-Nagel). Quantification of the isolated DNAs was performed using NanoDrop 8000 Spectrophotometer (Thermo Fisher Scientific) and Qubit 2.0 Fluorometer (Invitrogen, Life Technologies).

SNVs chosen as genetic markers to build the genotyping chip accomplished the following criteria. We included 42 CS-SNVs and 6 Sh-SNVs, only heterozygous alternative variants were selected from the deep-filtered list, based on their ability to discriminate among the four resequenced clones. For the CS-SNVs, equivalent number of variants for each clone were chosen. Sh-SNVs were picked for their ability to differentiate between the resequenced clones with a long history of clonal propagation in Argentina, from those with a short or null history in this country. Since we were particularly interested in identifying genetic markers that could consistently resemble that historical aspect across our samples. We also intended the chosen SNVs to be distributed across different chromosomes to better represent the genome-wide diversity. In total, 48 sequences containing one SNVs of interest (Supplementary Table S3) were provided to the Genomics Service Sequencing and Genotyping Unit (UPV/EHU) (Bizkaia, Spain) to design probes for a Fluidigm chip (https://www.fluidigm.com/) and perform the genotyping. Each experiment allowed to simultaneously genotype 48 samples using 48 SNVs, in a two steps reaction. In the first step, the target region containing the position to be genotyped is amplified using two pre-amplification primers (locus-specific primer and specific target amplification). In the second step, an additional PCR amplifies a portion of that target SNVs region, using the locus-specific primer and two fluorescently labeled allele-specific primers, which are internal primers containing either the first or the second allele respectively. Finally, the genotype is determined by measuring the fluorescence intensity of both alleles using the Fluidigm genotyping analysis software.

### d. Inter-clonal genetic diversity analyses

To assess the degree of genetic variation among ‘Malbec’ clones, biallelic genotypes were coded to sequences in the *fasta* format. In first place, we performed a phylogenetic analysis with MEGA v7.0.26 [^61^] including only the four resequenced clones and using PN40024 genotype as outgroup. A Neighbor-Joining tree was estimated based on the deep-filtered list of SNVs, using uncorrected *p*-distances and nodes’ support were obtained after 200 bootstrap iterations.

In second place, we performed a Median-Joining Network [^62^] analysis with POPART software [^63^]; to screen the diversity across all the genotyped samples, identify the number of different genotypes, their frequencies and phylogenetic relations. Afterwards, we obtained a single representative sequence for each of the identified genotypes to reconstruct a Neighbor-Joining tree with MEGA v7.0.26 [^61^], using the same parameters described above. Genetic diversity was also analyzed considering each SNVs position as an independent marker, by estimating codominant-genotypic distances among the identified genotypes with GenAlEx v6.5 [^64^]. Genetic distances among genotypes were analyzed with a model-free approach of Principal Coordinate Analysis (PCoA), to detect potential groups of genotypes closer related among each other. We estimated the proportion of the total molecular variance that is explained by the variance between groups through an AMOVA, we obtained the *Phi*PT parameter recommended for distances obtained from codominant genotypic data, *p-*value was obtained after 900 bootstrap iterations. Both PCoA and AMOVA were also performed with GenAlEx v6.5.

## Acknowledgments

This work was supported by Agencia Nacional de Promoción Científica y Tecnológica (ANPCyT): PICT2015-0822, MINCyT/ANPCyT (FONTAR)-CDTI ‘IBEROGEN’; CONICET (Bilateral PCB-II, CONICET–CSIC; MINECO BIO2017-86375-R). COST Action CA17111. We thank to M. Victoria Bertoldi for the assistance with laboratory work at IBAM-CONICET.

## Authors contributions

L.C. wrote the manuscript and performed: samples collection, laboratory work for library preparation and SNVs validation; bioinformatic and statistical analyses on the final set of genomic data. N.M. conducted bioinformatic analyses of raw data processing, variant calling, filtering and SNVs classification. C.M. collaborated with samples collection and laboratory work. P.C.B. provided insight on the SNPs and SNVs variant calling analysis and filtering. L.B. and C.S. collaborated with samples collection at Mercier nursery and provided information about historical background of the analyzed accessions. S.G.T. collaborated with samples collection at INTA. C.R. contributed to obtain and genotype samples from Europe. J.I. and J.M.M.Z provided insight on the design of the genotyping experiments. D.L. designed and coordinated the entire project. All authors carefully read and helped to improve the final content of the manuscript.

## Additional information

### Data access

SRA files containing raw genomic data for the four resequenced ‘Malbec’ clones is available at NCBI, BioProject: PRJNAXXXXXX.

### Supplementary information

Figures and tables complementing this paper are available in separate files

