## Supplemental figures for "Clonal propagation history shapes the intra-cultivar genetic diversity in ‘Malbec’ grapevines"

Figure S1. Example of four SNVs validated through Sanger sequencing. One clone specific SNV for each of the four re-sequenced clones are displayed. In each case the electropherograms show that the position of interest (indicated with the blue line) is heterozygous for the expected clone (on top) and homozygous as the reference in the other three clones, as expected.

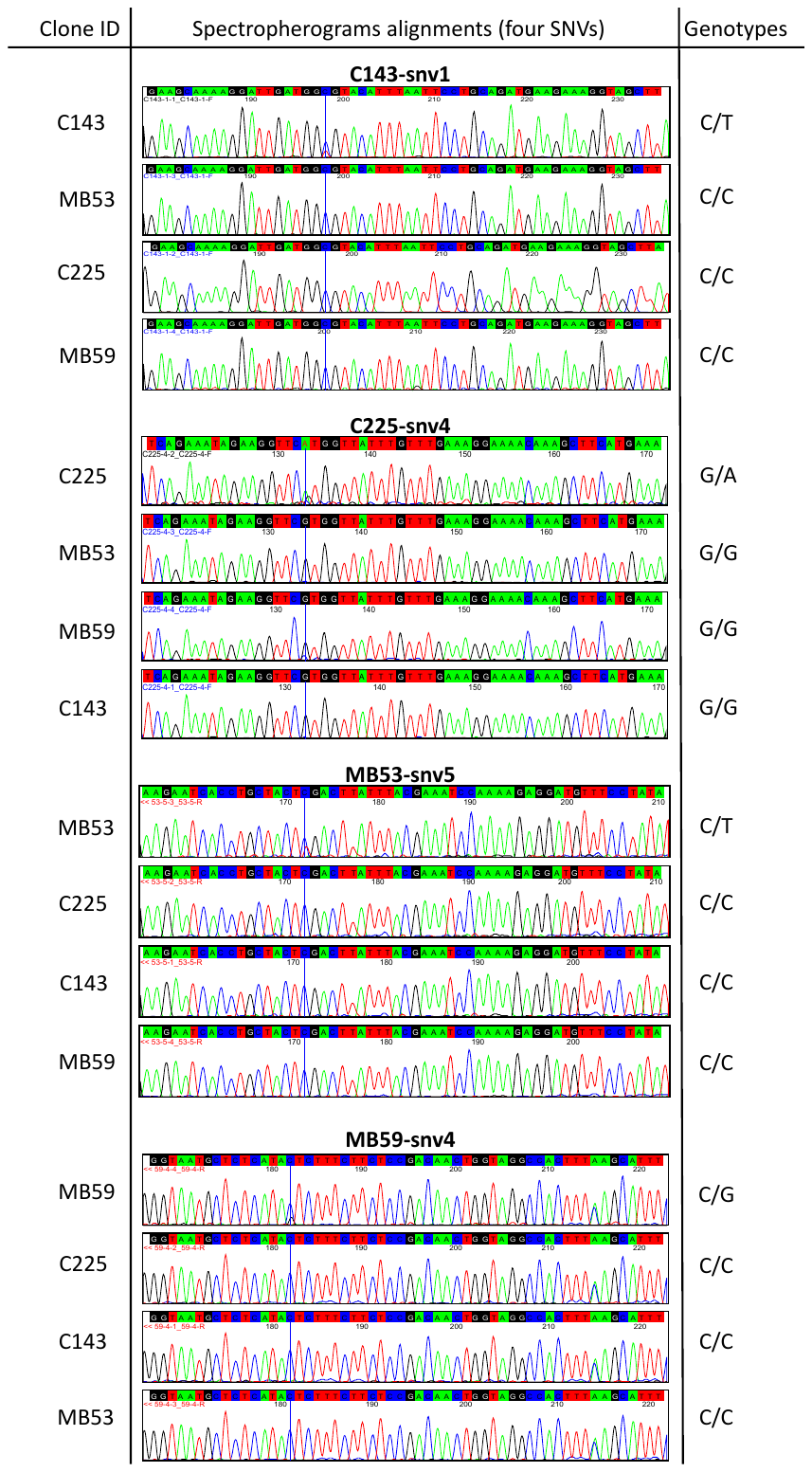

Figure S2. Genotypes network based on the 22 SNVs with more than two samples showing an alternative allele. We recovered the main nine groups of genotypes than the analysis using the 41 SNVs, but only two singleton genotypes: C225 and MB59.

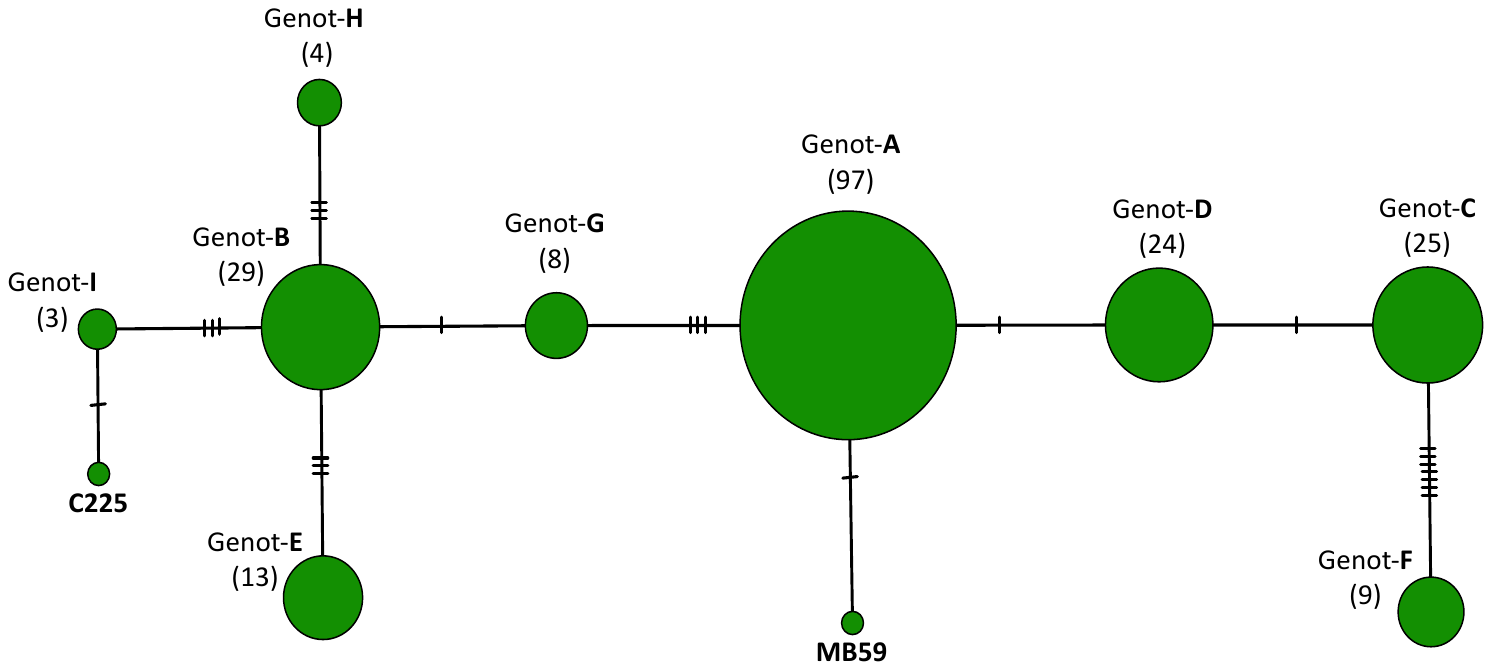

Figure S3. Principal coordinates analysis based only on the four shared SNV (Sh-SNV). PCoA recovers the distinction between Groups Ar and Fr.

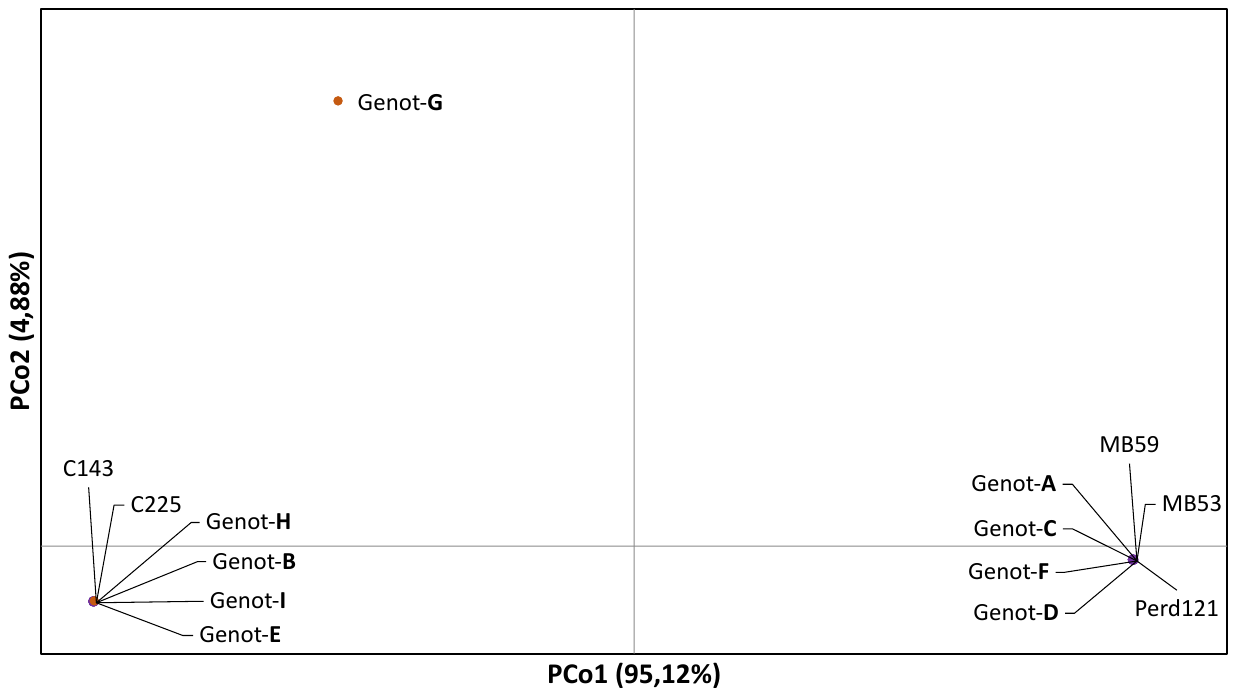
