## Supplemental Table S1 for "Clonal propagation history shapes the intra-cultivar genetic diversity in ‘Malbec’ grapevines"

**Table S1**. Raw data of paired-end (PE) reads obtained for each resequenced clone. Alignment stats values reported after filtering, percentage and coverage of reads alignment to *Vitis vinifera* L. reference genome (PN40024).

|  | Raw data | | | Alignment stats | |
| --- | --- | --- | --- | --- | --- |
| Clone ID | PE reads obtained | Reads size | Total sequence (Gb) | Covered % | Depth coverage (+SD) |
| C143 | 86,859,172 | 125 | 10.85 | 77.55 | 28.47 (70.66) |
| C225 | 94,388,045 | 125 | 11.79 | 77.9 | 30.26 (59.52) |
| MB53 | 90,054,921 | 125 | 11.25 | 78.01 | 31.29 (48.7) |
| MB59 | 90,125,307 | 125 | 11.26 | 77.94 | 29.96 (40.61) |
