## Supplemental Table S4 for "Clonal propagation history shapes the intra-cultivar genetic diversity in ‘Malbec’ grapevines"

**Table S4**. Details of the 41 SNVs employed in the final analyses. The name of the SNV refers to the re-sequenced clone for which it was identified. Genomic position is relative to PN40024, as well as the reference allele (Ref) state. The alternative allele (Alt) was observed in that position only for one of the four clones. Finally, the genotypic frequencies for each SNV based on the 214 successfully genotyped samples.

|  |  | Allelic state | | Genotypic frequencies | | |
| --- | --- | --- | --- | --- | --- | --- |
| SNVs-ID | Genomic position | Ref | Alt | XX | XY | YY |
| C143-snv1 | chr10:2146290 | C | T | 213 | 1 | 0 |
| C143-snv2 | chr10:2250888 | C | T | 201 | 13 | 0 |
| C143-snv3 | chr11:2533840 | T | C | 213 | 1 | 0 |
| C143-snv4 | chr12:5997741 | T | C | 213 | 1 | 0 |
| C143-snv6 | chr14:12612349 | C | G | 201 | 13 | 0 |
| C143-snv7 | chr16:18527058 | T | C | 213 | 1 | 0 |
| C143-snv11 | chr7:5723085 | G | A | 201 | 13 | 0 |
| C143-snv13 | chrUn:8135037 | C | T | 213 | 1 | 0 |
| C143-snv14 | chrUn:30681019 | G | A | 213 | 1 | 0 |
| C225-snv1 | chr11:13104147 | G | A | 213 | 1 | 0 |
| C225-snv4 | chr15:1260006 | G | A | 209 | 1 | 4 |
| C225-snv6 | chr18:18692633 | T | C | 213 | 1 | 0 |
| C225-snv8 | chr19:15624611 | G | T | 213 | 1 | 0 |
| C225-snv9 | chr4:17654346 | G | A | 210 | 4 | 0 |
| C225-snv11 | chr6:10576840 | G | A | 210 | 4 | 0 |
| C225-snv15 | chrUn:1879577 | C | T | 210 | 4 | 0 |
| MB53-snv1 | chr1:8947277 | G | A | 205 | 9 | 0 |
| MB53-snv2 | chr12:11248831 | C | T | 213 | 1 | 0 |
| MB53-snv3 | chr17:8192917 | G | A | 213 | 1 | 0 |
| MB53-snv4 | chr17:14078937 | G | A | 180 | 34 | 0 |
| MB53-snv5 | chr18:2701946 | C | T | 205 | 9 | 0 |
| MB53-snv6 | chr19:9264852 | C | T | 205 | 9 | 0 |
| MB53-snv7 | chr4:18276730 | A | C | 180 | 34 | 0 |
| MB53-snv8 | chr6:16432984 | A | G | 205 | 9 | 0 |
| MB53-snv9 | chr7:10631864 | C | T | 205 | 9 | 0 |
| MB53-snv10 | chr7:16870565 | C | G | 205 | 9 | 0 |
| MB53-snv11 | chrUn:1602277 | G | A | 205 | 9 | 0 |
| MB59-snv1 | chr1:4231365 | G | A | 213 | 1 | 0 |
| MB59-snv2 | chr10:9009503 | C | A | 209 | 5 | 0 |
| MB59-snv4 | chr14:612099 | C | G | 213 | 1 | 0 |
| MB59-snv5 | chr14:25250166 | C | T | 213 | 1 | 0 |
| MB59-snv9 | chr17:8058225 | C | G | 212 | 2 | 0 |
| MB59-snv10 | chr18:9067411 | T | C | 213 | 1 | 0 |
| MB59-snv11 | chr19:15786950 | C | T | 213 | 1 | 0 |
| MB59-snv12 | chr4:8993951 | A | G | 213 | 1 | 0 |
| MB59-snv14 | chr9:5534019 | T | C | 213 | 1 | 0 |
| MB59-snv15 | chr9:18487093 | T | C | 213 | 1 | 0 |
| MB53-MB59-snv2 | chr17:161911 | C | T | 50 | 164 | 0 |
| MB53-MB59-snv3 | chr19:22916577 | C | T | 58 | 156 | 0 |
| MB53-MB59-snv5 | chr5:22788880 | C | G | 58 | 156 | 0 |
| MB53-MB59-snv6 | chrUn:14723119 | C | T | 58 | 156 | 0 |
