## Supplemental Table S7 for "Clonal propagation history shapes the intra-cultivar genetic diversity in ‘Malbec’ grapevines"

Table S7. Brief description of the five ‘Malbec’ mass selections from Mercier Argentina nursery sampled.

| **Mass selection ID** | **Vineyard origin** | **Plants selection criteria** | **Selection outcome** |
| --- | --- | --- | --- |
| Perdriel | District: Perdriel  Department: Luján de Cuyo  Province: Mendoza | Absence of virus and fungus. Absence of *millerandage.*  Loose cluster, medium size grape, red rachis at maturity. | Average Productivity: 155 qq/Ha  Average cluster weight: 120 gr  Average total polyphenols index (IPT) = 60. |
| Vista Flores | District: Vista Flores  Department: Tunuyán  Province: Mendoza | Absence of virus and fungus. The complete vineyard was preserved because of high. quality wines production. | Average productivity: 170 qq/Ha  Average cluster weight: 190 gr  Average total polyphenols index (IPT) = 87. |
| Las Compuertas | District: Las Compuertas  Department: Luján de Cuyo  Province: Mendoza | Absence of virus and fungus.  Absence of *millerandage.*  Low productivity and high-quality wine production. | Average productivity: 110 qq/Ha  Average cluster weight: 90 gr  Average total polyphenols index (IPT) = 50. |
| Las Paredes | District: Las Paredes  Department: San Rafael  Province: Mendoza | Absence of virus and fungus.  Medium size grape and cluster, red rachis at maturity. | Average productivity: 193 qq/Ha  Average cluster weight: 110 gr  Average total polyphenols index (IPT) = 45. |
| Cobos | District: Agrelo  Department: Lujan de Cuyo  Province: Mendoza | Absence of virus and fungus.  Grapes skin showing intense purple coloration. | Highly variable productivity: ranging between 160 and 317 qq/Ha.  Average cluster weight: 150 gr  Average total polyphenols index (IPT) = Not Available |
